# Integrated Cross-Species Translation and Biophysical Multi-Scale Modeling Links Molecular Signatures and Locomotory Phenotypes in Spaceflight-Induced Sarcopenia

**DOI:** 10.1101/2025.05.03.652040

**Authors:** Brendan K. Ball, Hammad F. Khan, Jee Hyun Park, Krishna Jayant, Deva D. Chan, Douglas K. Brubaker

## Abstract

Age-related skeletal muscle deterioration, referred to as sarcopenia, poses significant risks to astronaut health and mission success during spaceflight, yet its multisystem drivers remain poorly understood. While terrestrial sarcopenia manifests gradually through aging, spaceflight induces analogous musculoskeletal decline within weeks, providing an accelerated model to study conserved atrophy mechanisms. Here, we introduced an integrative framework combining cross-species genetic analysis with physiological modeling to understand mechanistic pathways in space-induced sarcopenia. By analyzing rodent and human datasets, we identified conserved molecular pathways underlying microgravity-induced muscle atrophy, revealing shared regulators of neuromuscular signaling including pathways related to neurotransmitter release and regulation, mitochondrial function, and synaptic integration. Building upon these molecular insights, we developed a physiologically grounded central pattern generator model that reproduced spaceflight-induced locomotion deficits in mice. This multi-scale approach established mechanistic connections between transcriptional changes and impaired movement kinetics while identifying potential therapeutic targets applicable to both spaceflight and terrestrial aging-related muscle loss.

## INTRODUCTION

Space exploration is rapidly developing with the increasing number of missions to the International Space Station and beyond. As humans push towards longer spaceflights and journeys to other planets, understanding the physiological changes due to space travel has become critical. Despite decades of active research, our understanding of the biological burden of spaceflight remains limited. Moreover, this limitation is further exacerbated by the effects of prolonged exposure to the microgravity environment^1^ and cosmic radiation,^2,3^ which has been proven to be detrimental to human health.^4^

A well-documented consequence of space exploration is the deterioration of musculoskeletal health. Astronauts who return from space experience muscle atrophy, impaired muscle performance, and loss of strength.^5,6^ This process of muscle degradation resembles sarcopenia, a progressive age-related condition that often emerges over several decades and leads to muscle loss and function.^7^ While this condition progresses over an individual’s lifetime on Earth, the development of sarcopenia-like pathology is accelerated under microgravity conditions.^8^ Despite noted observations of muscle and strength breakdown,^9^ the exact mechanisms and pathways by which space-induced sarcopenia progresses are poorly understood.

There are major challenges to understanding how conditions in space affect human health. First, there are limited experimental samples and astronauts that have been studied on the International Space Station, often due to space constraints, costs, and accessibility.^10^ This is important because studying physiological changes in space-like conditions on Earth does not effectively translate to the harsh conditions in spaceflight. Additionally, astronauts on average spend 6 months per space mission tour, and thus there is limited information on the longer-term effects of space travel and sample size to study such effects.^11^ To address this challenge, previous studies aimed to improve our understanding of spaceflight and sarcopenia through the use of *in vitro*^12^ and *in vivo* rodent models.^13^ Despite these efforts, these approaches did not account for the heterogeneity of the human population and the biological differences across mammalian species. As a result, our understanding of a mechanistic pathway to sarcopenia conditions is largely unknown.

To overcome this limitation, we leveraged a framework known as Translatable Components Regression (TransComp-R), a computational model that synthesizes multi-omics data from preclinical rodent models and human data to predict disease outcomes in humans.^14,15^ The TransComp-R pipeline has been successfully applied to overcome other biomedical challenges including Alzheimer’s disease,^16–20^ osteoarthritis,^1^ and inflammatory bowel disease.^14,21^ We coupled this approach with a biophysically grounded model of central pattern generator (CPG) networks,^22,23^ therefore enabling multiscale interrogation of sarcopenia.

In this study, we used TransComp-R to show that transcriptomics data from mouse models under spaceflight conditions predicts human sarcopenia outcomes. We performed cross-species computational modeling to synthesize spaceflight mouse and sarcopenia human data to identify potential biological pathways that are involved in muscle degeneration. To link the observed space-related sarcopenia to neuromuscular function, we built a network of leaky integrate-and-fire neurons to build a computational CPG model that recapitulated locomotion kinetics in healthy conditions. Importantly, incorporation of the predictive transcriptomic data into the model generated similar phenotypic gait responses observed under sarcopenia conditions. Thus, this approach enables an improved understanding of mechanisms associated with space-induced sarcopenia and may inform therapeutic avenues for sarcopenia and other muscle-deteriorating conditions related to spaceflight and on Earth.

## RESULTS

### Mouse spaceflight principal components separate human sarcopenia and control conditions

We accessed RNA-sequencing spaceflight mouse data from the NASA Open Science Data Repository (OSDR). The publicly available spaceflight dataset (OSD-103) contained 16-week-old C57BL/6J male mice with a sample size of n=6 ground controls and n=6 spaceflight conditions.^24,25^ The spaceflight group was housed at the International Space Station for 37 days before euthanasia and collection of quadricep muscle samples. The ground control group was housed on Earth for reference. We then acquired RNA-sequencing data of patients with sarcopenia from Gene Expression Omnibus (GEO). We selected three separate cohort studies that studied sarcopenia among different racial demographics, including Caucasian (GSE111006), Afro-Caribbean (GSE111010), and Chinese (GSE111016) descent.^26,27^ Our integration of multiple cohorts allowed us to reference our findings across different human backgrounds.

We applied the TransComp-R model to identify principal components (PCs) that encoded transcriptomics variation of spaceflight mice that predicted sarcopenia outcomes in humans. The TransComp-R pipeline works by computing a mouse principal component analysis (PCA) space and projecting human data into space. Mouse PCs that encode transcriptomics capable of stratifying human disease from control groups are then identified by a regression model and can be biologically interpreted through pathway enrichment analyses and follow-up studies.

With the TransComp-R model, we first matched homologous gene pairs across the mouse and all three human datasets (**Figure 1a**). We next applied principal component analysis (PCA) on the mouse dataset to identify PCs that explained 80% of the cumulative variation and projected the human dataset with the Chinese cohort with sarcopenia and control conditions into the mouse PC space.

**Figure 1.**
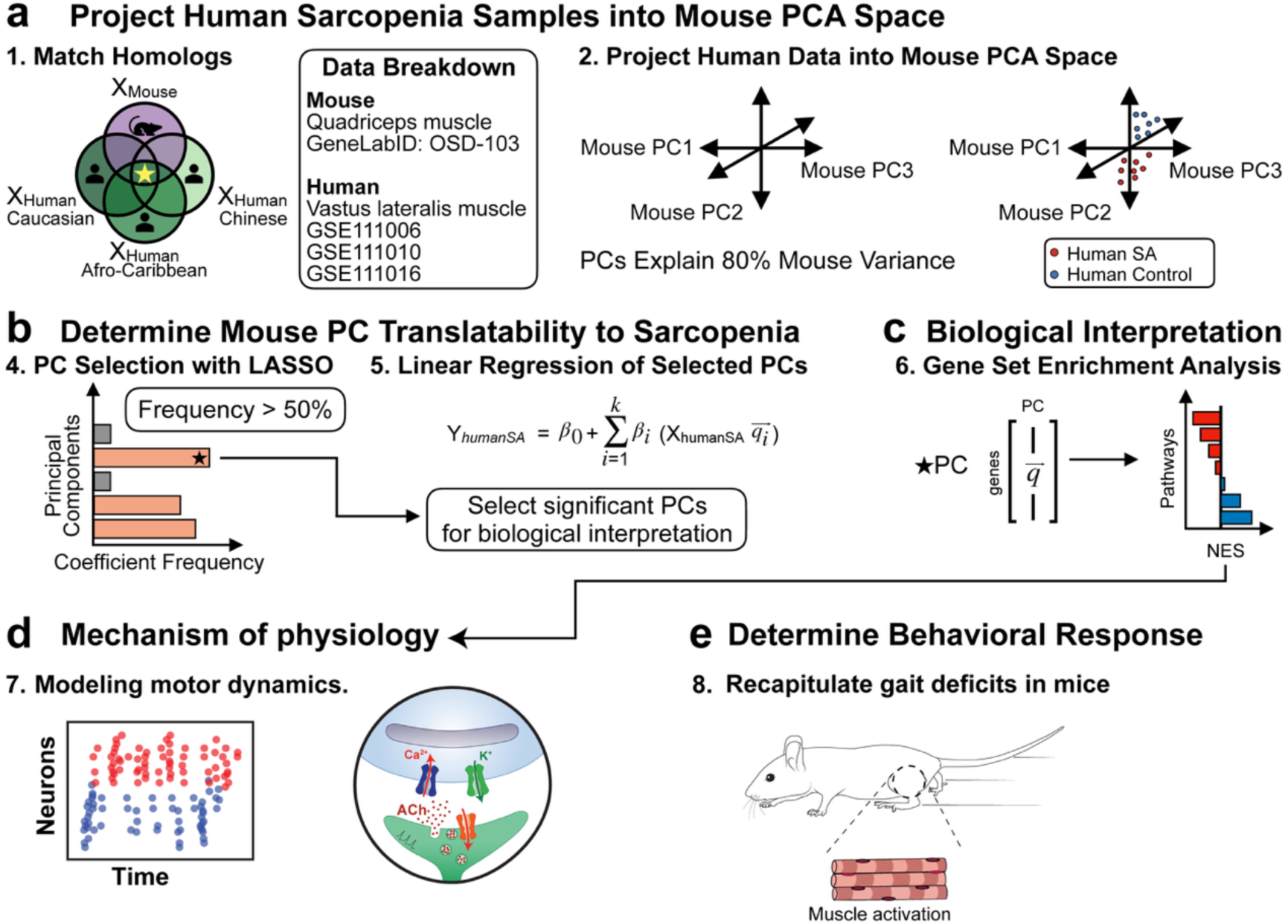
Mouse to human modeling workflow. **(a)** Homologous pairs between mouse and human are first matched. Human sarcopenia and control groups (Chinese cohort) are projected into the space flight mouse PCs. The human Caucasian and Afro-Caribbean cohorts were used for downstream validation analysis. **(b)** The PCs selected by LASSO are regressed against human sarcopenia and control outcomes to determine significance. **(c)** Significant PCs from the generalized linear model are used for downstream biological analysis. (**d**) Biophysical modeling of motor dynamics. (**e**) Recapitulate behavior deficits in space-induced sarcopenia based on genetic determinants.

After synthesizing the mouse and human transcriptomics data together, we selected PCs that were relevant in predicting human sarcopenia condition with LASSO (least absolute shrinkage and selection operator) (**Figure 1b**). The selected PCs from LASSO were used in a generalized linear regression against human sarcopenia and control groups. The significant PCs were then interpreted using gene set enrichment analysis (GSEA). Lastly, we validated our findings with the Caucasian and Afro-Caribbean cohorts that also contained sarcopenia and control data.

After matching for homologous pairs, we identified 12,484 genes shared across the mouse and human datasets. We constructed a mouse PCA space and projected the Chinese cohort and identified 7 PCs that explained 80% of the variance (83.4% var. exp). We next calculated how much variance the mouse PCs explained in the human data to understand the potential of translation from mouse to human (**Figure 2a**). Among the seven mouse PCs, we found PC1, PC3, and PC6 explained a higher ratio of human variance compared to mouse variance. This finding suggests that PC1, PC3, and PC6 may have a more pronounced ability to capture variance across species than other PCs.

**Figure 2.**
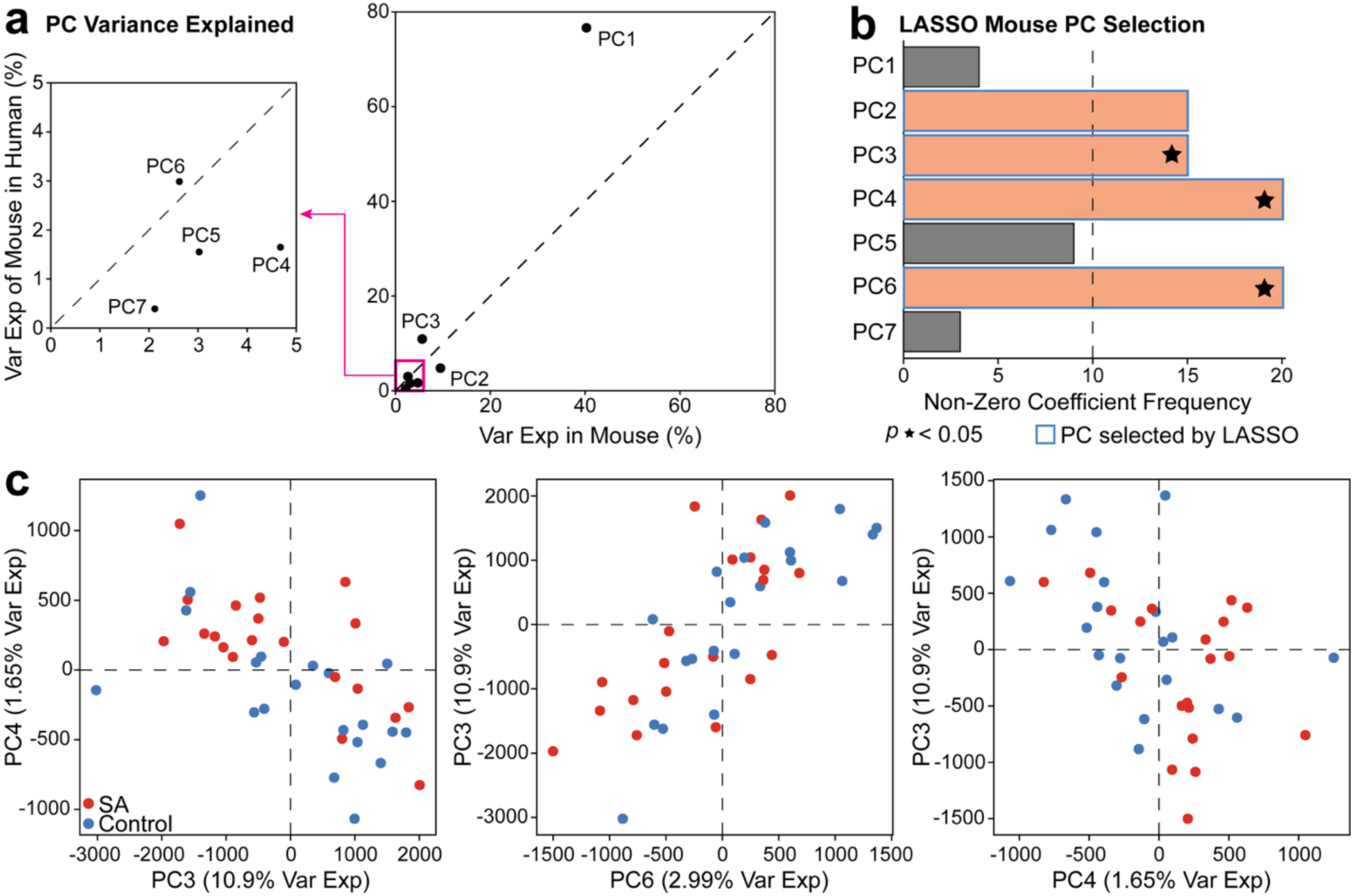
Significant mouse PCs distinguish sarcopenia and control conditions in humans. **(a)** The percent variance explained of mouse PCs and how much they explain in human data. **(b)** PC selection using the LASSO. PCs selected by LASSO that were significant in a GLM (p value < 0.05) are denoted with a star. **(c)** Plots of significant mouse PCs separating sarcopenia and control in humans (SA=sarcopenia).

Of the seven PCs, we identified PC2, PC3, PC4, and PC7 with our LASSO model (selected more than half of the twenty random rounds) to be relevant eigenvectors with power to stratify human sarcopenia and control patients (**Figure 2b**). We created a generalized linear regression model incorporating these four PCs and found PC3, PC4, and PC6 to be significantly separating the sarcopenia and control groups (p = 0.0286, 0.0371, and 0.0253, respectively). We next visualized the three significant PCs and found that the transcriptomics variation encoded on each PC separated the human sarcopenia and control groups (**Figure 2c**).

### Metabolic and immune-associated pathways on cross-species principal components distinguish patients with sarcopenia

Having identified significant murine PCs that distinguish between human sarcopenia conditions, we conducted GSEA on the PC loadings and identified pathways related to neuroactive ligand receptor interactions, immune pathways, and cellular structures enriched for human sarcopenia on mouse spaceflight PC3 (**Figure 3a**). Genes that were involved with neurodegenerative diseases and metabolic pathways were associated with healthy human groups. This indicates that these genes may become disrupted under sarcopenia conditions and encode human phenotype predictive cross-species biological signatures.

**Figure 3.**
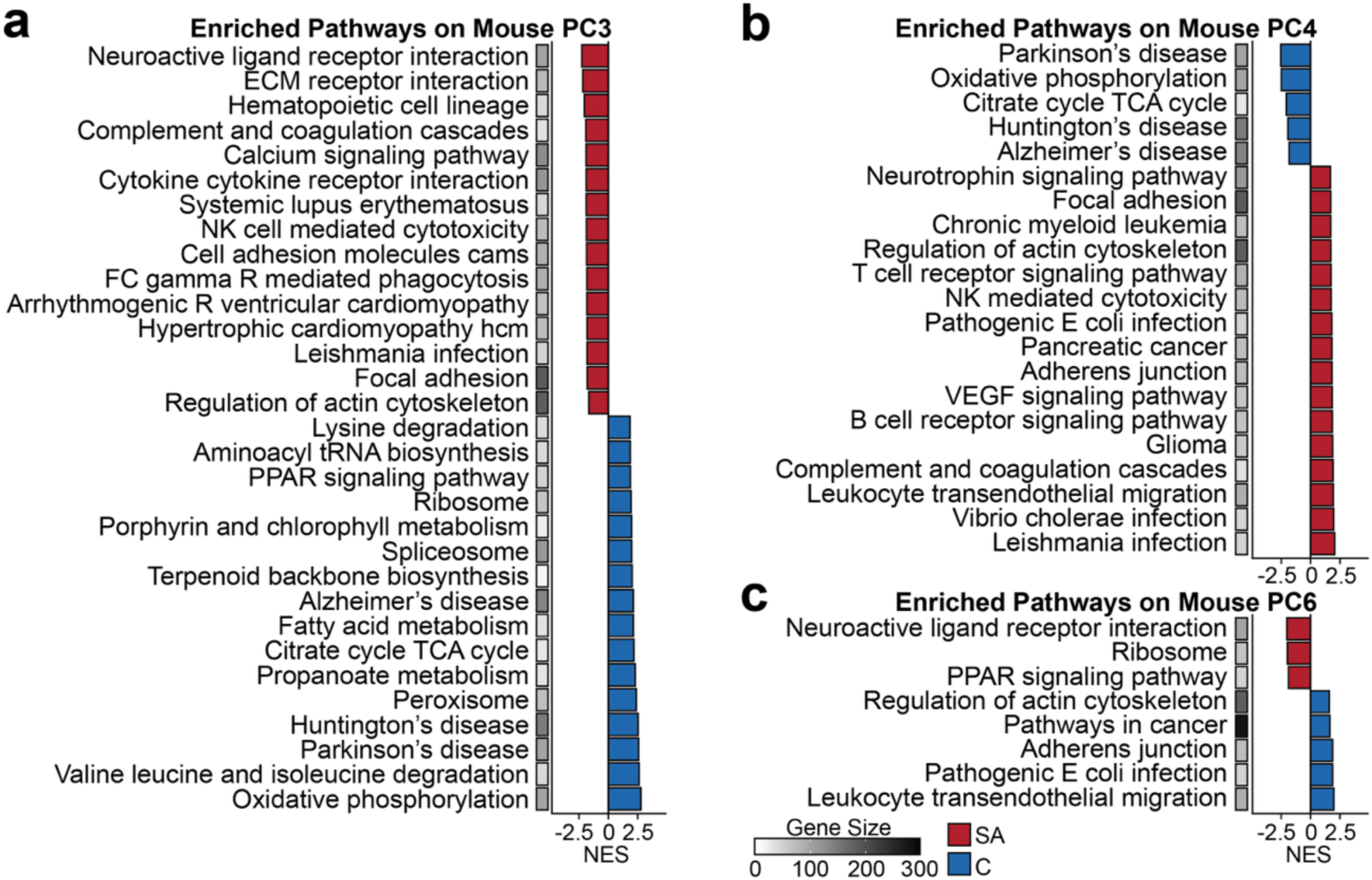
KEGG pathway enrichment analysis. Biological pathways enriched on **(a)** PC3, **(b)** PC4, and **(c)** PC6. All pathways are significant with a Benjamini-Hochberg adjusted p<0.01. The gene size represents the number of genes enriched in the biological pathway. Pathways are shown with associations to human sarcopenia (red) and control (blue) based on the directionality of separation for each PC. The bar plots are represented by their respective normalized enrichment score (NES). (SA=sarcopenia, C=control).

On the mouse spaceflight PC4, we found pathways involved with immune signaling and cellular structure to be enriched in the sarcopenia group (**Figure 3b**). Among the pathways enriched on PC6, we found neuroactive ligand receptor interaction and PPAR signaling associated with sarcopenia, whereas regulation of actin cytoskeleton, adherens junction, and leukocyte transendothelial migration were enriched in control (**Figure 3c**).

We also performed GSEA using the hallmark curation and found signaling mechanisms involved with the immune system to be involved with sarcopenia (**Supplementary Figure 1**). Additionally, we found oxidative phosphorylation to be consistently enriched for the control group in PC3, PC4, and PC6. Such results further support our findings with the reported KEGG pathway analysis. These findings also show that genes contributing to the immune responses, as well as mitochondrial function associated with oxidative phosphorylation may play important roles in the muscle degenerative process associated with sarcopenia in humans and can be measured in a predictive capacity in murine spaceflight models.

### Significant PC gene loadings from the Chinese cohort are translatable to Caucasian and Afro-Caribbean demographics

After identifying biological pathways associated with sarcopenia and control, we assessed whether our findings generalized across human demographic groups. We constructed three PLS-DA models using combined Caucasian and Afro-Caribbean human sarcopenia data and genes extracted from each significant mouse spaceflight PC (PC3, PC4, PC6) that our TransComp-R model as having cross-species predictive value in the Chinese cohort. Our objective was to see whether the biological signatures identified in the significant cross-species PC’s of TransComp-R could generalize to other demographic groups.

We combined the Caucasian and Afro-Caribbean sarcopenia datasets to determine if findings from these populations were also reflected by our findings from the Chinese cohort. The Caucasian and Afro-Caribbean datasets contained further annotations on muscle weakness in the subjects beyond those reported in the Chinese cohort. We first excluded control patients with any reported history of muscle weakness or muscle loss to avoid confounders. We combined the Caucasian and Afro-Caribbean datasets model and verified that batch effects were not present with PCA (**Supplementary Figure 2**). Breaking down the combined and processed demographics, we analyzed n=28 healthy Caucasians (Age: 72.6±2.3 standard deviations), n=4 sarcopenia Caucasians (Age: 74.3±3.6), n=14 Afro-Caribbean (76.4±7.4), and n=9 sarcopenia Afro-Caribbean (76.8±7.9) individuals.

We selected a panel of genes for each model from each TransComp-R identified PC, selecting the top 10 and bottom 10 genes ranked by their murine PC loadings, for PLS-DA predicting human sarcopenia vs. control groups in the combined Caucasian and Afro-Caribbean dataset. Genes strongly contributing to the model contained a calculated variable importance of projection (VIP) score greater than 1.

We demonstrated a separation of control and sarcopenia with the combined human dataset (**Figure 4a**). With murine spaceflight PC3 genes, the human samples were separated among the first latent variable (LV). On this LV, we found genes that are strongly associated with sarcopenia, which included *DLG3*, *CALB2*, *LHFPL6*, and *ADAM33* (**Figure 4b**). The *DLG3* and *CALB2* genes are associated with neuron signaling, whereas *LHFPL6* and *ADAM33* are involved with cell-to-cell signaling and communication. Oppositely, genes such as *PPP1R12B* and *CNTF are associated* with the control groups. These genes are involved in helping muscle contraction through myosin phosphatase activity and enhancing survival factors for neuronal cells, respectively.

**Figure 4.**
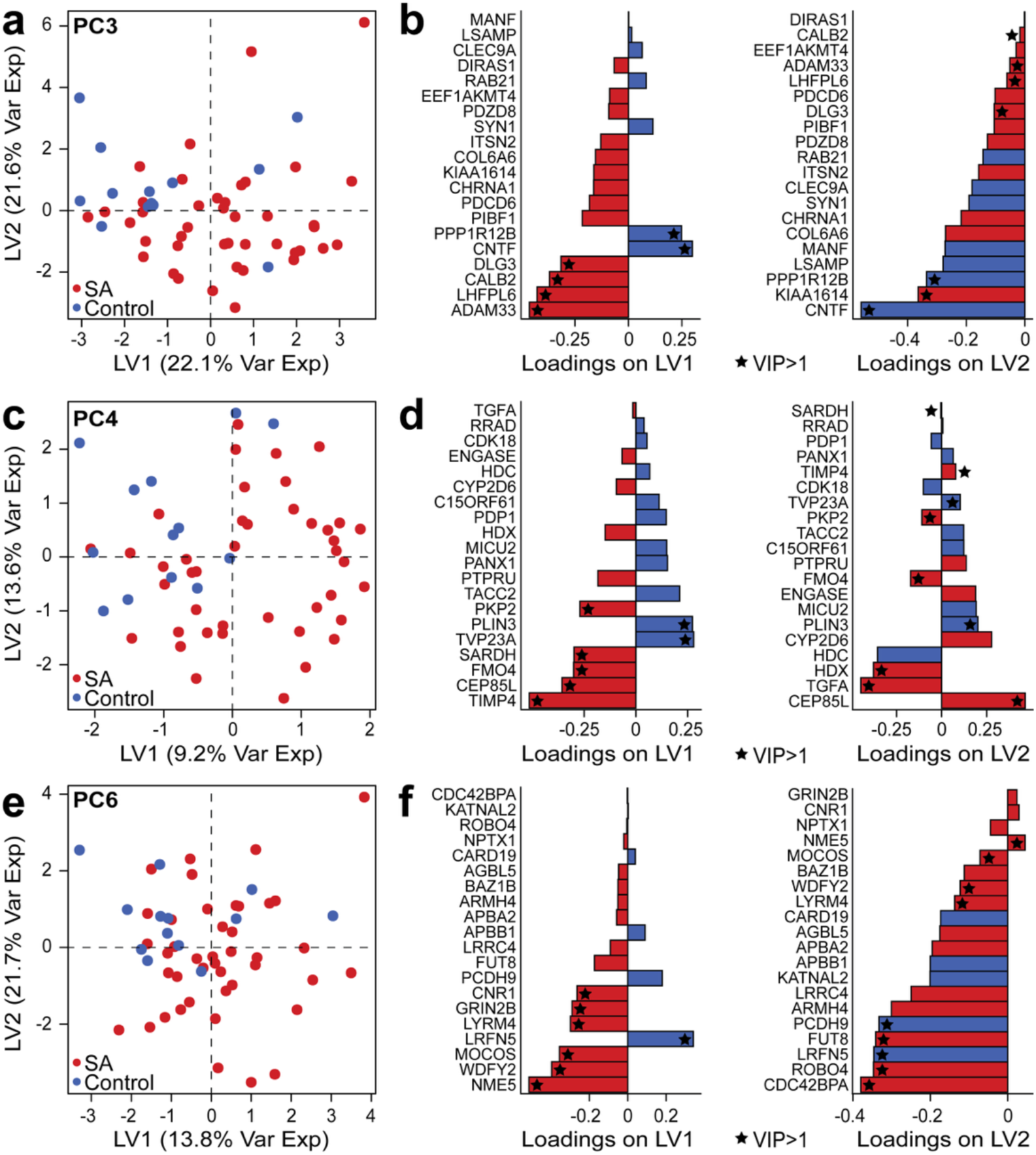
Gene expression driving separation among Chinese sarcopenia cohorts is also reflected in Caucasian and Afro-Caribbean populations. **(a)** PLS-DA on the top 10 and bottom 10 genes identified in the spaceflight PC3. **(b)** The loadings of the first two latent variables for PC3. **(c)** PLS-DA for PC4. **(d)** Loadings for PC3. **(e)** PLS-DA for PC6. **(f)** Loadings for PC6. Loadings with a VIP>1 are labeled with a star. The color on the loadings represents the highest contribution to sarcopenia or control by the specific gene. (SA=sarcopenia).

We also performed PLS-DA on mouse spaceflight PC4 genes and demonstrated that LV1 discriminated between the control and sarcopenia human groups (**Figure 4c**). We identified *PKP2*, *SARDH*, *FMO4*, *CEP85L*, and *TIMP4* genes to associate with sarcopenia, whereas *PLIN3* and *TVP23A* were associated with control (**Figure 4d**). These genes have known associations with cellular structure, metabolism, and vesicle transport. For murine spaceflight PC6, we similarly found the control group clustered on the negative score of LV1, and the sarcopenia group clustered among positive scores (**Figure 4e**). This separation was defined by *CNR1*, *GRIN2B*, *LYRM4*, *MOCOS*, *WDFY2*, and *NME5* for sarcopenia. For the control groups, we found *LRFN5* as the only gene contributing to the PLS-DA separation with a VIP>1. Interestingly, the *CNR1* and *GRIN2B* genes are linked to neurotransmission and brain function whereas *LYRM4* and *MOCOS* are connected to mitochondrial metabolism.^28–30^ The *LRFN5* gene we found associated with the control group is associated with cell-to-cell communication among neuronal cells.

### In silico drug screening analysis identifies candidate potential therapeutics for preventing spaceflight-associated sarcopenia

After confirming these mouse PCs are reflective in sarcopenia across multiple human demographics, we leveraged a computational correlation analysis to shed light on potential therapeutics associated with space-induced sarcopenia. We used the Library of Integrated Network-Based Cellular Signatures (LINCS) Consensus Signatures that contains characteristic direction coefficients based on gene expression profiles of 33,609 drugs.^31^ We analyzed 2,558 of these drugs after excluding duplicates and those that did not contain any reported cellular targets. We used a Spearman’s correlation analysis between the characteristic direction coefficient values of affected genes from a respective drug to the loadings on selected mouse PCs. Our goal was to determine medications that may have potential therapeutic benefits by identifying drugs that induced gene signature changes opposite to the transcriptomics profile of spaceflight-induced sarcopenia. Here, we focused on significant drugs that were approved by the Food and Drug Administration (FDA) to ensure our findings were met with standards for safety and effectiveness.

We found 135 drugs significantly correlated with the gene loadings encoded on the mouse PC3 (**Figure 5a**). We consulted the 135 drugs to the FDA Orange Book (March 2025 version) and determined 37 FDA-approved drugs. From the selected FDA-approved drugs, isradipine had the highest correlation coefficient of 0.2246 and is used to treat hypertension through calcium channel blockers. We also found mestinon (ρ=0.2184), a medication primarily used for muscle weakness condition in myasthenia gravis, and sunitinib (ρ=0.2003), an anti-cancer medication, as drugs associated with reversing the sarcopenia gene signatures. Among the drugs with the highest association of inducing sarcopenia-like gene expressions, we found two medications used to treat schizophrenia (ariprazole, ρ=-0.1958, and lurasidone, ρ=-0.1689), and temsirolimus (ρ=-0.1756), used for renal cell carcinoma.

**Figure 5.**
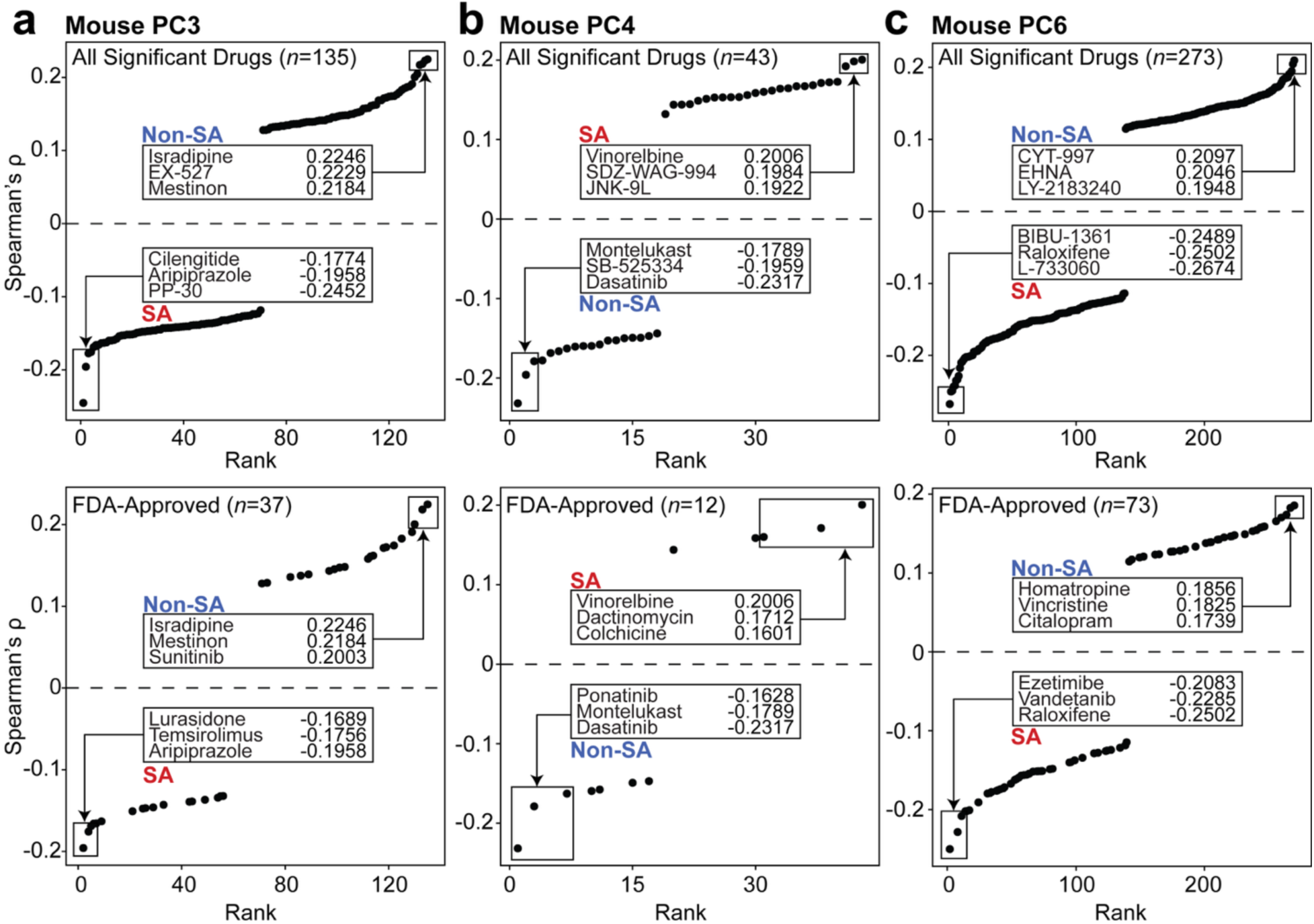
Gene expression correlational analysis for therapeutic discovery. All significant drugs and a subset of significant drugs selected by their FDA-approval status for **(a)** mouse PC3, **(b)** mouse PC4, and **(c)** mouse PC6. The top and bottom three medications ranked by their Spearman’s ρ value are labeled by sarcopenia (signatures correlated to genes elevated in sarcopenia) or non-sarcopenia (signatures correlated to genes elevated in healthy conditions). (SA=sarcopenia, Non-SA=healthy condition).

On mouse PC4, we identified 43 significant drugs associated with spaceflight induced sarcopenia gene expression, 12 of which were FDA-approved (**Figure 5b**). Dasatinib, a chemotherapy drug used to treat cases of leukemia, had the most negative correlation score (ρ=-0.2317). Other medications that correlated with the non-sarcopenia profile included montelukast (ρ=-0.1789, treats asthma symptoms) and ponatinib (ρ=-0.1628, chronic myeloid leukemia treatment). The FDA-approved medications that most associated with sarcopenia-like conditions included vinorelbine (chemotherapy for non-small cell lung cancer), dactinomycin (chemotherapy medication), and colchicine (anti-inflammatory to treat gout attacks).

With mouse PC6, we identified 273 significant drugs, 73 of which were FDA-approved. Homatropine (ρ=0.1856), vincristine (ρ=0.1825), and citalopram (ρ=0.1739) were among the drugs with the highest correlation to reversing spaceflight-induced sarcopenia gene expression profiles. These medications are used for muscarinic acetylcholine receptor antagonists, leukemia and cancers, and selective serotonin reuptake inhibitor antidepressants, respectively. Conversely, medications with the highest associations to inducing a spaceflight-induced sarcopenia disease signatures included raloxifene (ρ=-0.2502), vandetanib (ρ=-0.2285), and ezetimibe (ρ=-0.2083). Raloxifene is used for individuals undergoing menopause to address osteoporosis and breast cancer risk reduction. Vandetanib and ezetimibe are used to treat medullary thyroid cancer and to reduce high cholesterol levels in the blood, respectively.

### Multivariate analysis reveals rodent gait deficits related to orbital spaceflight

Space-induced muscle atrophy represents a compelling model for investigating fundamental mechanisms of sarcopenia, just as sarcopenia is a compelling disease model for studying ways to prevent space-induced muscle atrophy. We demonstrated with pathway enrichment analysis that neuroactive ligand receptor interaction and PPAR signaling were associated with sarcopenia in both human and rodent models, suggesting conserved genetic determinants for sarcopenia.^32^ To further interrogate the relationship between sarcopenia changes at the molecular level, we performed multivariate analysis of locomotory deficits in rodents following prolonged microgravity exposure aboard the International Space Station.

We accessed the spaceflight behavioral assay available on the NASA database (OSD-478), which contains a prospective collection of gait data between mice undergoing spaceflight in orbit aboard the ISS and ground control over 35 days.^33^ Pre-flight and post-flight gait measurements were conducted across a total of n=40 mice (n=20 mice per group), measuring multiple aspects of locomotion and gait kinematics. We conducted an unsupervised analysis of effects on gait dynamics across spaceflight and ground control by projecting the data into a mouse PC subspace to extract behavior dynamics that were most affected during spaceflight.

We used PCA of gait dynamics to reveal distinct behavioral signatures between pre- and post-flight conditions (**Figure 6a, b**). We clustered the treatment groups based on chi-squared confidence ellipse and noted a distinct separation in PC space that was more pronounced in spaceflight groups compared to ground controls, suggesting specific microgravity-induced adaptations across mice as opposed to temporal artifacts. The first PC captured 25.5% of total variance followed by 14.8%, and 11.3%, for PC 2 and 3, respectively. Approximately 80% of locomotory variation was explained by just eight components (**Figure 6c**), suggesting strongly shared effects across all mice undergoing spaceflight.

**Figure 6.**
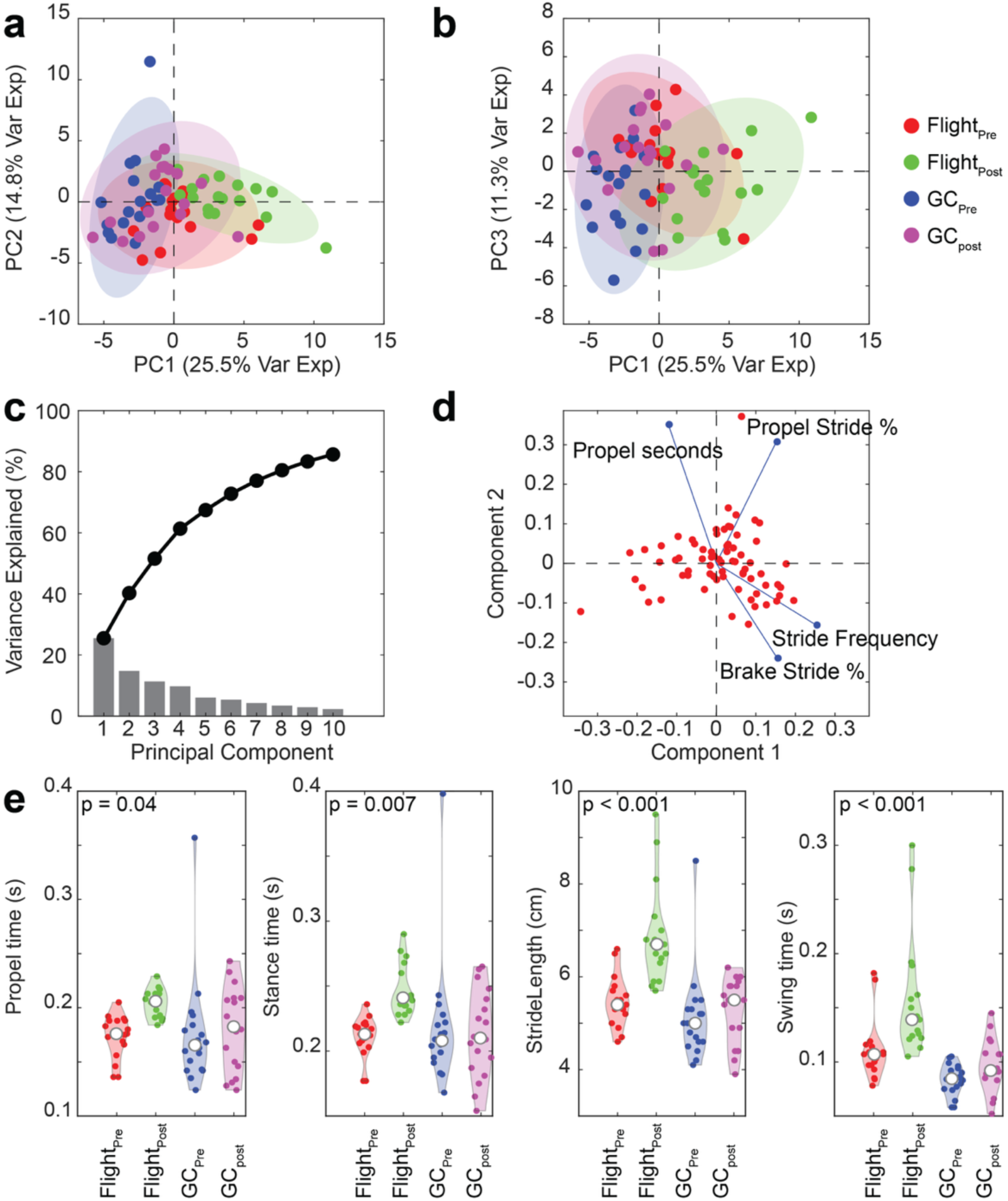
Behavioral alterations during spaceflight. **(a)** PC projection of the first and second component axis with behavioral variables under pre- and post-flight conditions with ground controls. **(b)** PC projection of the first and third PC components. **(c)** Scree plot of variance explained for the first 10 PC components. **(d)** A bi-polar plot of the top 4 variable behaviors among the flight and ground control groups. **(e)** Highest weighted behavioral alteration properties determined by PC analysis (n = 40 mice total, one-way ANOVA test).

To investigate behavioral contributions, we conducted a loading analysis of tracked gait properties during locomotion and identified key gait parameters contribution to PC variance in the publicly available mouse data (**Figure 6d**). For example, among the first two components, we identified propulsion parameters and stride as primary contributors to observed variance (**Figure 6d**). We found similar results when conducting systematic testing across all gait parameters across mice (**Supplementary Figure 3**). To further quantify this, we selected the most contributing gait parameters across the first 4 PC components and performed inferential statistics across mice. We found significant alterations in propel time (p=0.04), stance time (p=0.007), stride length (p<0.001), and swing time (p<0.001) following orbital habitation (**Figure 6e**). These deficits suggest impairments in both stride generation and balance feedback mechanisms essential for coordinated movement.^34^ Similar observations were noted in humans undergoing functional gait tests.^35^ Moreover, this analysis demonstrated clear locomotor deficits in mice following spaceflight exposure. Notably, the propelling force and stride kinematics were strongly altered compared to ground control. These coordinated changes could suggest disruption in multiple neural pathways governing both balance maintenance and gait kinetics, consistent with functional adaptations to microgravity conditions.^35,36^

### Physiological modeling of gait characteristics

To understand the physiological mechanisms underlying space-induced locomotory deficits and key molecular determinants of sarcopenia, we established a biologically plausible physiological model that captured the essential dynamics of spinal locomotor circuits under normal terrestrial conditions. This computational framework serves as a substrate to later assess microgravity-induced alterations, demonstrating how changes in biomolecular network activity directly translate to muscle function and ultimately to observable gait parameters.

Our network model consisted of leaky integrate-and-fire neurons following well-established membrane potential and gating dynamics.^22,23,37–39^ To capture oscillatory dynamics often found during locomotion, we assigned distinct flexor and extensor neural populations that were inhibitory connected to one another. This enabled a subset of neurons to generate patterned activity in the absence of input to both regions, thus reflecting CPG dynamics (**Figure 7a**).^37^ Simulation of the population model drove characteristic oscillatory patterns in membrane potential dynamics between flexor and extensor circuits under normal gravity conditions (**Figure 7b**), with flexors exhibiting more rapid depolarization, consistent with their role in initiating limb movement. The computational network successfully reproduced rhythmic activity reflecting central pattern generators, with flexor neurons (red) showing synchronized burst patterns distinct from extensor neurons (blue) during simulated normal locomotion (**Figure 7c**). We used the cumulative output of each group to infer muscle activation patterns that were used to extract gait dynamics (**Figure 7d**). Gait kinematics exhibited a strong phase-coherent response (67%) between extensor and flexor periods (**Figure 7e**) reflecting coordination between muscle groups.

**Figure 7.**
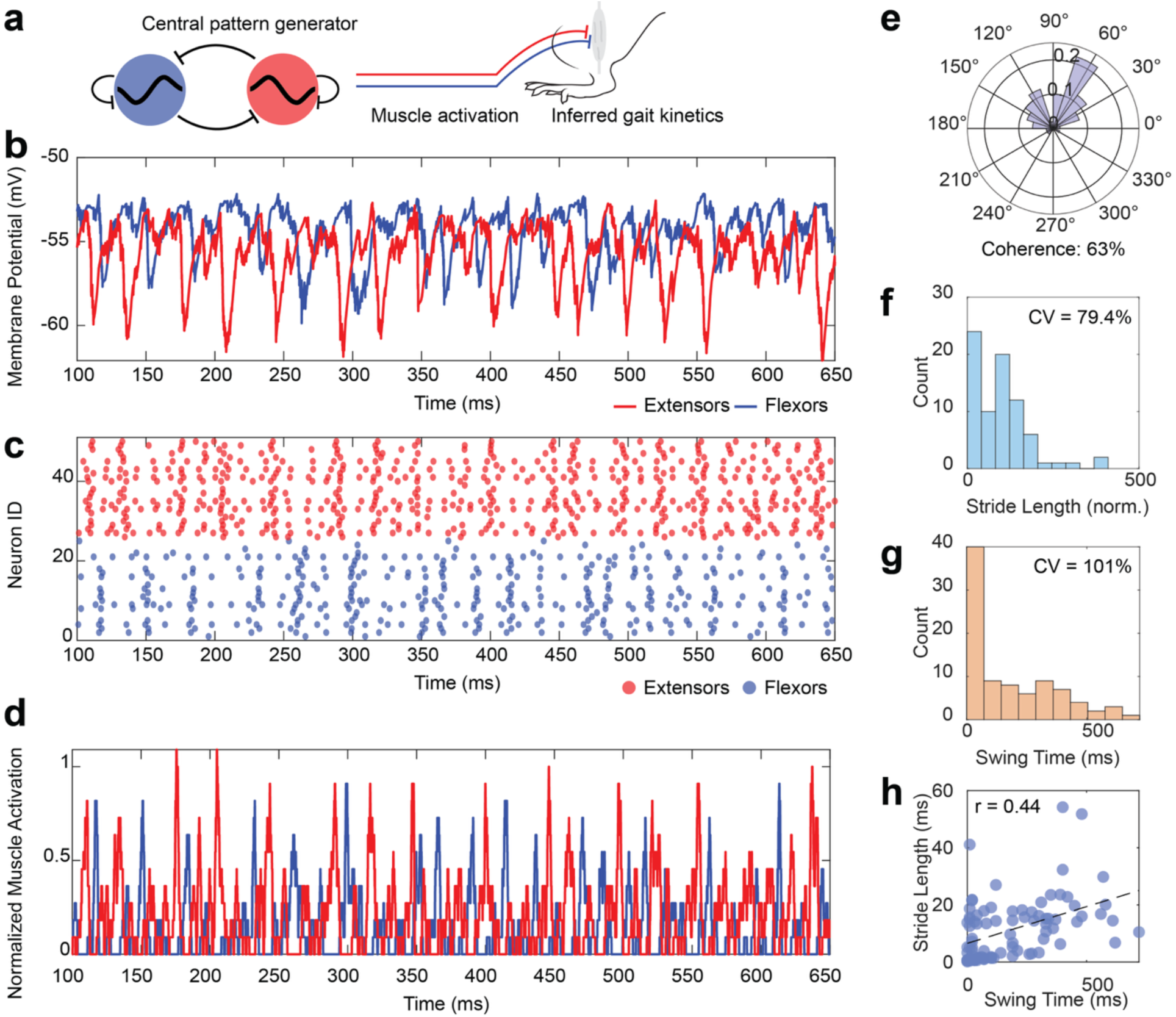
Motor neuron population model for locomotory behavior. **(a)** Schematic of the mechanistic physiological model. **(b)** Averaged population membrane dynamics from a leaky integrate-and-fire model. **(c)** Rastergram of 50 neurons spiking during constant input. Neurons are colored based on assigned flexor or extensor dynamics. **(d)** Inferred muscle activity patterns as a function of population output. **(e)** Polar plot of phase relationship between flexor and extensor population dynamics. **(f)** Distribution of stride length with coefficient of variation. **(g)** Distribution of swing times with coefficient of variation. **(h)** Regression of correlation swing time and stride length.

Thus, this computational approach creates a powerful platform connecting three levels of analysis: neuronal firing patterns, muscle activation dynamics, and resultant gait kinetics (**Figure 7f-h**). The normal phase relationships, stride distributions, and swing patterns established here provide quantifiable baselines against which space-induced neural adaptations can be measured in subsequent experiments. By bridging cellular neurophysiology to whole-organism locomotion, this model enables mechanistic investigations into how microgravity disrupts motor control at multiple biological levels.

### Computational framework linking biomolecular determinants of sarcopenia to altered gait characteristics

To bridge the gap between the identified biomolecular determinants found by TransComp-R’s gene enrichment analysis and the physiology of gait characteristics, we modified our network model to exhibit sarcopenia phenotypic behavior. This was primarily informed by the identified ligand-gated genetic disruption (G*RIN2B* and *CNR1*), during spaceflight in both mice and human cohorts (**Figure 8a**). We modified the synaptic transmission probability and conductance, while biasing the total input variability, into the network of individual neurons in both flexor and extensor populations **(Figure 8b**). We noted that, upon this disruption, an observed reduction in phase-coherence between flexor and extensor dynamics compared to the healthy terrestrial network. (Coherence_healthy_ = 55%, Coherence_sarcopenia_ = 33%, **Figure 8c**). The change in network activity also drove a significant increase in both inferred gait swing time and stride length (p<0.0001, **Figure 8d, e**), most likely due to increased variability in the CPG network. Thus, these results recapitulate the same gait alterations observed in cohorts of mice that spend a significant time in space flight (**Figure 6e**).

**Figure 8.**
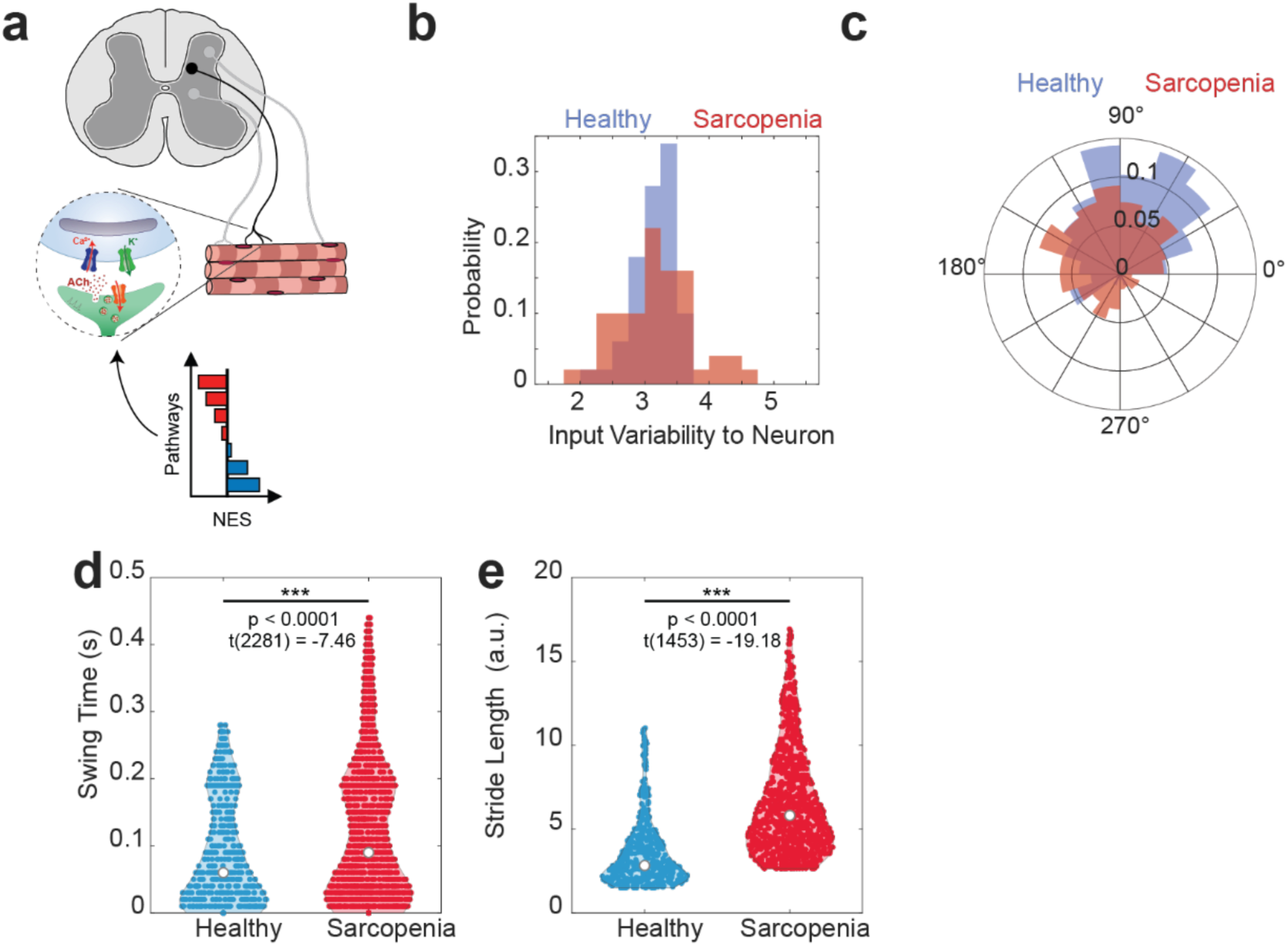
The physiological model illustrates a link between biomolecular determinants and excitation-inhibition balance during space-related sarcopenia. **(a)** Schematic showing a physiological model of the motor neuron to muscle activation alongside sarcopenia-related gene regulation from normalized enrichment score (NES). **(b)** Probability distribution of modeled input variability to neurons. **(c)** Healthy and sarcopenia population-related phase activity across neurons during inferred muscle activity. Note reduced phase strength during movement. Reported phase coherence for healthy and sarcopenia activity is R_healthy_ = 0.55 and R_sarcopenia_ = 0.33, respectively. **(d)** Modeled swing times of population response under healthy and sarcopenia (p = 0.00, t(1423) = -19.13, Student’s t-test). **(e)** Modeled swing times of population response under healthy and sarcopenia for calculated stride length (p = 0.00, t(2281) = -7.16, Student’s t-test).

This computational framework provided mechanistic insights connecting space-induced alterations in neural excitation/inhibition balance to the observed locomotor deficits. Our findings suggest that space-induced sarcopenia involves not only peripheral muscle atrophy but also fundamental adaptations in neural control systems, including altered central pattern generator function and disrupted coordination between antagonistic muscle groups. These neural adaptations likely contribute significantly to the gait abnormalities documented in our behavioral analysis and highlight potential targets for countermeasure development.

## DISCUSSION

Space-induced sarcopenia is a major physiological change that affects astronauts. We used transcriptomics muscle biopsies of mice exposed to microgravity and human sarcopenia data to understand the effects of space travel on humans. In this work, we presented a comprehensive framework to understand the biological pathway and its effect on locomotion phenotypes in space-induced sarcopenia. By leveraging TransComp-R, we used a mouse-to-human modeling approach to identify key biological pathways related to neurotransmitter release and regulation, synaptic integration, and mitochondrial dynamics associated with sarcopenia conditions. Likewise, a comprehensive analysis of locomotion behavior in mice exposed to spaceflight showed strong alteration in gait parameters which we mechanistically linked to a CPG network model, incorporating altered identified molecular pathways associated with sarcopenia conditions. Our results established cross-species predictive relationships for studying spaceflight-induced sarcopenia in a framework that links biomolecular determinants of sarcopenia to phenotypically resulting locomotion behavior.

We identified several immune system pathways altered under conditions of sarcopenia, which has also been observed under spaceflight conditions.^40–42^ A study reported immune dysregulation and the redevelopment of herpes virus among astronauts on a 6-month orbital space mission.^43^ A separate 6-month spaceflight that studied two crew members identified elevated proinflammatory biomarkers in muscle biopsy post-landing.^44^ During spaceflight, blood-derived cytokines including interleukins and tumor necrosis factor-alpha were also significantly elevated during flight.^45^ Further investigation of such immune pathways may improve our understanding of spaceflight-induced sarcopenia.

Metabolic pathways are also actively involved in muscle health as demonstrated by protein alterations during spaceflight.^46^ Stein and colleagues reported that whole-body protein synthesis was reduced by 45% about preflight measurements.^47^ In the same study, post-flight measurements returned to 91% of the pre-flight levels, indicating a lack of complete recovery from space travel.^47^ Additionally, we identified genes that were strongly associated with sarcopenia through neuromuscular signaling. Neuromuscular junction aging has been proposed to be involved in sarcopenia pathogenesis.^48^ The age-related denervation at the neuromuscular junction occurs before muscle atrophy^49^ and may be involved in sarcopenia pathogenesis.^50^

We applied a high-throughput drug screening analysis and identified FDA-approved medications that induced gene expression changes potentially linked to protection against or reversing spaceflight-induced sarcopenia. Of the three mouse PCs, we found particular interest in the findings presented on the mouse PC3 due to its ability to explain more variance in human data, indicating the highest potential for mouse-to-human translation. We found a calcium channel blocker (isradipine) and a medication primarily used for myasthenia gravis (mestinon) among the FDA-approved drugs that associated with gene expression changes of a healthy condition. Others have reported that calcium channel blockers like amlodipine^51^ and nifedipine^52^ are able to attenuate contraction-induced muscle damage muscular atrophy, respectively. A separate study that studied the continuous infusion of mestinon in rats showed improvements to muscle force with treatment post-immobilization-induced muscle weakness.^53^ Identifying potential therapeutics may provide insight for future follow-up studies surrounding calcium channel blockers and muscle-strengthening medications.

The biomechanical alterations we noted in our study from the behavioral assay align with previously documented changes in movement atrophy.^54,55^ The propulsion and stride deficits likely reflect preferential atrophy of fast-twitch extensor muscles that generate locomotive force, mirroring the fiber-type selectivity observed in terrestrial sarcopenia but compressed into a remarkably accelerated timeframe (∼35 days).^36,56^ The concurrent changes in stride parameters suggest simultaneous adaptation of central pattern generators and sensorimotor integration circuits. It is of interest that several key protein pathways were also affected in both humans and mice which could explain some of the downstream locomotion phenotypes. For example, the presence of *DLG3* and *GRIN2B* suggests alterations of synaptic integration across neurons within a CPG network. Interestingly, *GRIN2B* was upregulated in the skeletal muscle of aging mice.^57^ Such consequences would be reflected in how neural populations rhythmically generated activity between flexor and extensor periods. Indeed, when altering synaptic integration properties along the network in our CPG model, primarily synaptic release probability and input gain, we noted a clear reduction in firing coherence between flexor and extensor population, which recapitulates the behavior changes we observed in the gait dynamics.

Although our study identified several pathways to improve our understanding of space-induced sarcopenia, there are limitations to this work. First, due to limited publicly available data on mice that have been exposed to spaceflight, the sample sizes are smaller. Additionally, our mouse and human datasets only contained information from males, which neglects potential biological outcomes in females. Additionally, we only analyzed physiological changes in the quadricep muscle and did not investigate the contribution of other limb muscles. Future studies that incorporate additional demographic and sex-based information of both males and females, with the inclusion of other muscle biopsies and relevant tissue will largely build upon the findings we reported.

Likewise, there are limitations to our computational models. TransComp-R selects homologous gene pairs across mouse and human datasets; thus, genes that do not fit this criterion are omitted. As a result, there is a possibility that these omitted genes may have also served an active role in the physiological changes to spaceflight-induced sarcopenia. Our CPG network model incorporated leaky integrate-and-fire neurons with reciprocal connections, synaptic probabilities, and membrane dynamics. However, additional models can be incorporated to further explore the mechanistic contribution of protein altered during sarcopenia. For example, modeling a biophysically realistic model of neurons with full ion channel distributions and gating dynamics could be more informative on how changes at the protein level affect neural dynamics and CPG activity.

For example, one can investigate how excessive synaptic release alters integration properties of neurons and their firing rate. Further experiments are needed to causally link the synaptic and protein interactions to behavioral deficits observed during spaceflight or using alternative models of age-related sarcopenia.

Our study investigated the effects of sarcopenia resulting from spaceflight travel. We used TransComp-R and identified spaceflight mouse PCs that could predict changes in human age-associated sarcopenia in muscle biopsies. Our findings were then validated by matching a panel of genetic markers from these PCs in other human cohorts across different populations. Lastly, our CPG neuron model, incorporated with changes in cross-species predictive biomarkers of sarcopenia, was able to recapitulate healthy and space-induced sarcopenia phenotypes in locomotion. Thus, our findings in this work pave a path to link molecular pathways to behavioral phenotypes of spaced-induced sarcopenia.

## MATERIALS AND METHODS

### Data selection

We selected publicly available transcriptomics data from the NASA OSDR and GEO database with the prerequisite that the data contained a balanced number of samples with controls and processed tissue samples from similar quadriceps muscle regions in both mouse and human.

The mouse dataset was selected from the NASA OSDR. We focused our selection criteria on transcriptomics data that contained muscle samples near the quadriceps. We identified a mouse study (OSD-103) that contained a spaceflight and ground control group. We next searched the GEO database and identified three human sarcopenia datasets with controls and biopsy samples, each containing transcriptomics data of biopsy samples from the vastus lateralis muscle. The datasets represent a demographic population around the world, including Caucasian (GSE111006), Chinese (GSE111016), and Afro-Caribbean (GSE111010).

### Pre-processing and data normalization

We downloaded human transcriptomics sarcopenia data from the GEO database using the GEO2R platform with Bioconductor tools in R (*GEOquery* ver. 2.70.0, *Biobase* ver. 2.62.0, and *limma* ver. 3.58.1). We log_2_ transformed the data and z-score normalized the data per gene. We matched the human and mouse datasets for homologous one-to-one gene pairing using the *orthogene* package (ver. 1.8.0). Genes that did not contain homologous pairing were excluded from the model and further downstream analysis.

### Mouse to human computational modeling

We performed TransComp-R by performing PCA on the mouse space flight with both flight and ground control groups. We selected mouse PCs that explained up to a total of 80% cumulative variance to prevent overfitting of the data and inclusion of noise. We projected the human dataset (Chinese cohort GSE111016) into the mouse PC space. The projection of the human data into the mouse PCA space is a matrix multiplication of *X^s^ ^x^ ^n^* and *Q^n^ ^x^ ^p^* such that *s* is the row of human patients, *n* is the homologous gene pairs shared across both mouse and human, and *p* is the number of PCs that explain 80% cumulative variance. The equation is also shown:

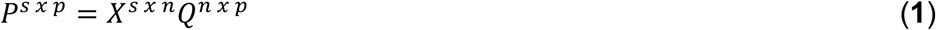

### Feature selection of mouse principal components

To select important features and regularize the mouse PCs that best predicted human sarcopenia condition, we performed LASSO across 20 rounds of 5-fold cross-validations by regressing the human sarcopenia and control group positions along the mouse PCA space against the human disease condition. Mouse PCs with a non-zero coefficient frequency greater than 10 of 20 rounds were selected for the generalized linear model. We performed the generalized linear regression using the equation:

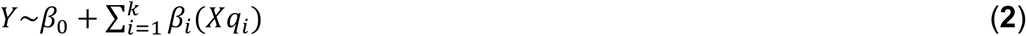

In the generalized linear model, the *Xq_i_* represents the projected human samples into mouse PCs that were selected by LASSO and *Y* is the human sarcopenia and control status. We selected mouse PCs with a p value less than 0.05 for downstream biological interpretation.

### Variance explained in human by mouse principal components

Using principal components encoding mouse variation and human data, we can calculate how much variation the mouse PCs explain the variation in humans. We calculated the variance explained by mouse PCs in human data with the equation:

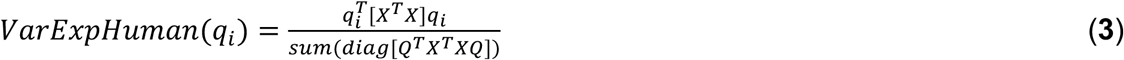

Where *X^s^ ^x^ ^n^* is a human data matrix represented by *s* number of subjects and *n* number of genes, and *Q^n^ ^x^ ^p^* is the mouse data represented by the loading coefficients of *n* number genes and *p* columns of principal components *q_i_*. The *T* represents the transpose of the data matrix.

### Gene set enrichment analysis

We selected PCs that significantly distinguished between human sarcopenia and control groups for downstream pathway analysis. To identify enriched pathways, we pre-ranked each significant PC by their respective loadings score and performed GSEA in R (*msigdbr* ver. 7.5.1, *fgsea* ver. 1.28.0, *clusterProfiler* ver. 4.10.1).^58–60^ We generated a holistic understanding of the enriched pathways by using the hallmark and KEGG curations from the Molecular Signatures Database. The parameters we used to identify enriched biological pathways included a minimum gene size of 5, a maximum gene size of 500, and an epsilon value of 0. We used the default parameter of 1000 permutations to determine the statistical significance. We defined biological pathways in both KEGG and hallmark to be significant if the Benjamini-Hochberg adjusted p value was less than 0.01.

### Gene selection for cross-population validation

To ensure our findings are reproducible in other datasets, we used the Caucasian and Afro-Caribbean datasets (GSE111006 and GSE111010) to determine if the findings in the Chinese cohort are reflective in other demographics. In both the Caucasian and Afro-Caribbean datasets, we first excluded control patients with a reported history of low muscle mass or low muscle strength and/or performance to ensure the control groups had no potential complications. We then concomitantly merged the two cohorts into a singular dataset and verified for batch effects visualized by PCA.

Using the Chinese cohort, we filtered for the top 10 and bottom 10 gene loadings from significant mouse PCs identified in our TransComp-R model to determine if transcriptomic variation encoded on these PCs could stratify sarcopenia and healthy groups in the Caucasian and Afro-Caribbean populations.

### Partial least squares discriminant analysis

We used the *mixOmics* package (ver. 6.26.0) to construct the PLS-DA models for each significant mouse PC that could significantly stratify human sarcopenia and control groups in the Chinese sarcopenia cohort.^61^

We identified the most important genes contributing to the model’s predictive ability by calculating the VIP scores. We used the *mixOmics* package to calculate the VIP scores for each given LV. The following equation:

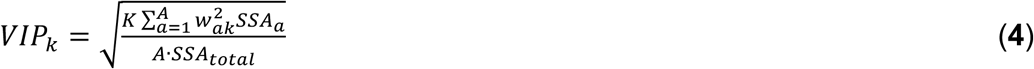

A given VIP score for a specific gene is calculated by *K* number of genes, *A* number of PLS-DA components, and *w_ak_* weight of the gene *k* in *a^th^*LV. *SSA_a_* and *SSA_total_* are the sum of squares in the *a^th^* LV and the total sum of squares in all LV components, respectively. We defined a VIP score greater than 1 to represent genes contributing above average to the model’s performance.

### In silico cross-species drug screening analysis

We computationally screened for drugs correlated with the mouse PCs predictive of sarcopenia for therapeutic potential. We used the L1000 Consensus Signatures Coefficient Tables (Level 5) from the LINCS database. Prior to data use, we referred to the LINCS small molecules metadata and removed all the drugs that did not report any known targets. To identify candidate drugs with potential therapeutic connections to sarcopenia, we integrated the differentially expressed genes (DEGs) from each drug in the LINCS database and the gene loadings from the mouse PCs predictive of human sarcopenia. The DEGs for each drug were selected by scaling the characteristic direction coefficients, the up- or down-regulation of a gene by the drug, to z-scores and obtaining genes, and their characteristic direction values, with p-value < 0.05.^31^

We paired the genes of the mouse PC loadings with the differentially expressed drug genes and performed Spearman’s correlation. We selected correlations that had a Benjamini-Hochberg adjusted *p* value less than 0.05. Based on the Spearman’s rho value, we ranked the drugs for each mouse PC that was processed.

### Multivariate analysis of locomotor behavior

We used the OSD-478 from the NASA OSDR for locomotor behavior analysis. Principal component analysis was applied to 47 locomotor behavioral parameters measured in n=35 C57BL/6J mice across four experimental groups: pre-flight spaceflight (pre-flight), post-flight spaceflight (post-flight), pre-flight ground control (GC pre), and post-flight ground control (GC post). Parameters included stride kinematics (length, duration, variability), limb coordination metrics, and ground reaction forces collected during treadmill locomotion.

### Behavior and spaceflight cluster validation

All behavioral and locomotion analysis was performed using MATLAB. Group separation was visualized in reduced-dimensional space by projecting z-scored data onto the first three PCs. We calculated 95% confidence ellipses for each experimental group using:

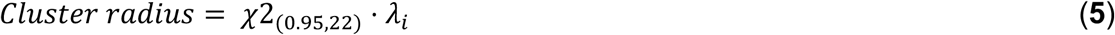

where χ=5.991 and *λ_i_* represents the eigenvalue of the *i^th^*PC. Multivariate analysis of variance (ANOVA) with Hotelling’s T² post-hoc tests assessed group differences in PC space (α=0.05, Bonferroni-corrected).

### Feature selection and hypothesis testing

Behavioral variables most associated with spaceflight effects were identified by examining PC loadings. For each of the first three PCs, we selected variables with absolute loadings exceeding the 95th percentile of loading magnitudes (equivalent to the top 5 variables per PC).

### Motor neuron population model for movement

To model motor neuron population for biophysical extraction of movement, we simulated a population of 50 leaky integrate-and-fire neurons with equal proportions of flexor and extensor connectivity to muscle output. Membrane potential dynamics followed biophysically realistic neurons (**Table 1**). Neuronal populations between extensor and flexors were reciprocally connected with inhibitory conductance, generating coupled generation of spiking activity resembling pattern generation circuits (CPG) for locomotion.

**Table 1.** Parameters for the CPG model.

| Variable | Value |
| --- | --- |
| Resting potential (mV) | -65 |
| Reset potential (mV) | -70 |
| Membrane resistance (MΩ) | 10 |
| Synaptic conductance (nS) | 0.4 |
| Inhibitory reversal potential (mV) | -80 |
| Synaptic decay (ms) | 5 |
| Mean membrane time constant (ms) | 10 |
| Refractory period (ms) | 3 |

We also incorporated “kick and decay” kinetics during reciprocal inputs of neuronal populations. Together the change in voltage for each neuron was characterized as:

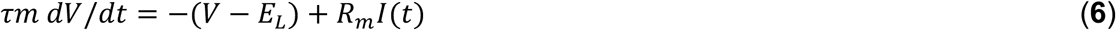

where τ*m* is the membrane time constant, V is the membrane potential, *E_L_* is the leak reversal potential, *R*_*m*_ is the membrane resistance and *I(t)* is the input current as a function of time.

### Inference of locomotory activity

Population firing rate activity was temporally smoothed with a 5 ms Gaussian kernel. Peak population activity bursts between flexor and extensor neurons were used to calculate the phase difference and cycle lengths which were used to infer stride length and variability, respectively. The correlation of variation was calculated by taking the mean and standard deviation of cycle lengths between population bursts. To infer swing times, the duration of peak responses in the flexor population was calculated and the differences were taken for each cycle. Similarly, the stance time was taken as the time from the end of the previous swing cycle and the next swing cycle. The ratio of both swing and stance times yielded the swing stance ratio.

## ACKNOWLEDGEMENTS

BKB and HFK is supported by the NSF GRFP (DGE-1842166). DDC is supported in part by the DARPA Young Faculty Award (Army Research Office Contract W911NF21103272) and National Science Foundation (Awards 1944394 and 2149946). DKB is supported by an award from the Good Ventures Foundation, Open Philanthropy, and start-up funds from Case Western Reserve University.

## AUTHOR CONTRIBUTIONS

BKB: Conceptualization, data curation, formal analysis, investigation, methodology, visualization, writing-original draft, writing-review & editing. HFK: Conceptualization, data curation, formal analysis, investigation, methodology, visualization, writing-original draft, writing-review & editing. JHP: Data curation, methodology, writing-review & editing. KJ: Project administration, resources, writing-review & editing. DDC: Conceptualization, methodology, project administration, resources, funding acquisition, writing-review & editing. DKB: Conceptualization, funding acquisition, methodology, project administration, resources, writing-review & editing.

## DATA AVAILABILITY

All human transcriptomics data used from the study is available through Gene Expression Omnibus with accession numbers GSE111006, GSE111010, and GSE111016. Mouse transcriptomics data is available through the NASA Open Science Data Repository with accession OSD-103. Mouse behavioral data is available with accession OSD-478 on the NASA OSDR.

## SUPPLEMENTARY INFORMATION AND CODE AVAILABILITY

All code used for the analysis is available at https://github.com/Brubaker-Lab/MouseFLTandHumanSA. All supplementary information including figures and data is also made available on the GitHub repository.

## SUPPLEMENTARY INFORMATION

**Supplementary Figure 1.**
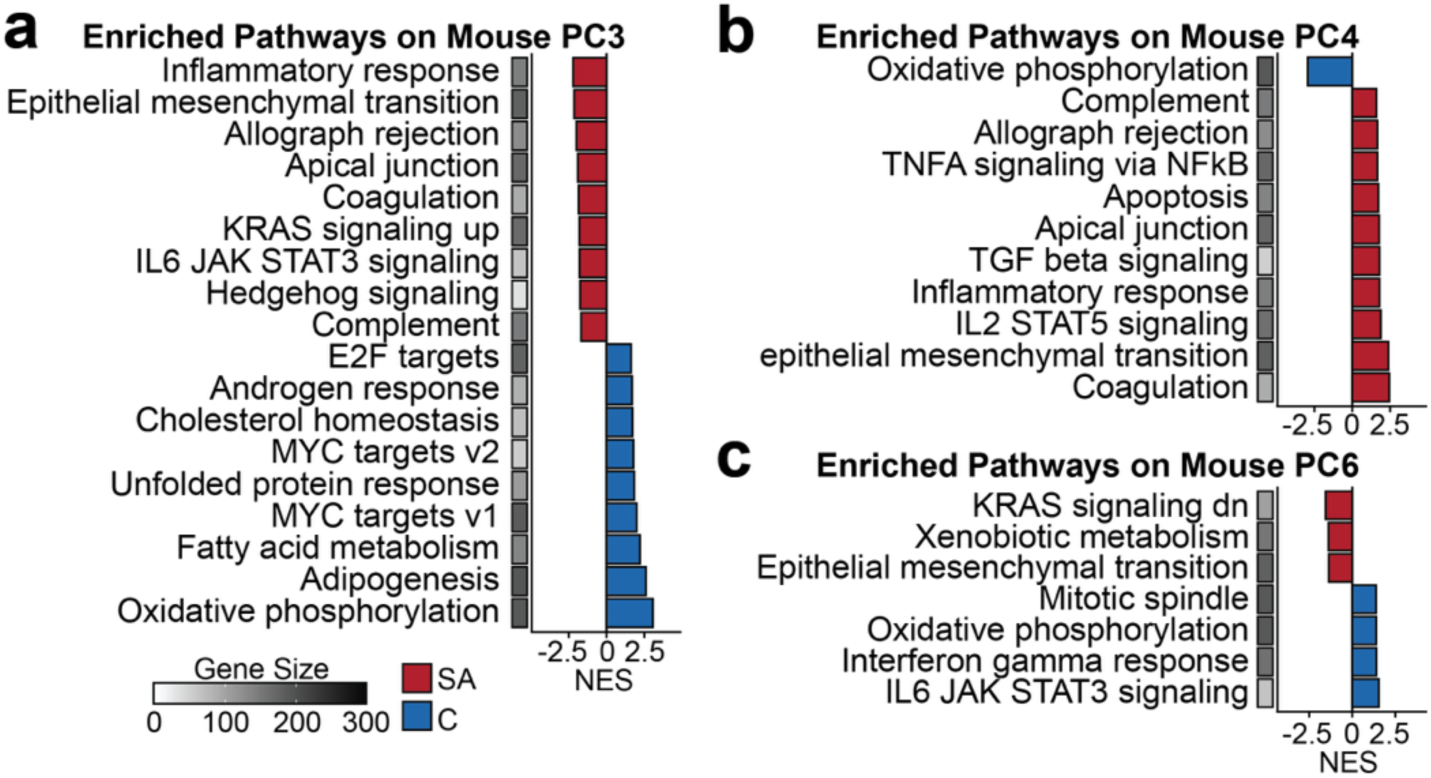
Hallmark pathway enrichment analysis. **(a)** Significantly enriched pathways on mouse PC3, **(b)** PC4, and **(c)** PC6. (SA=sarcopenia, C=control).

**Supplementary Figure 2.**
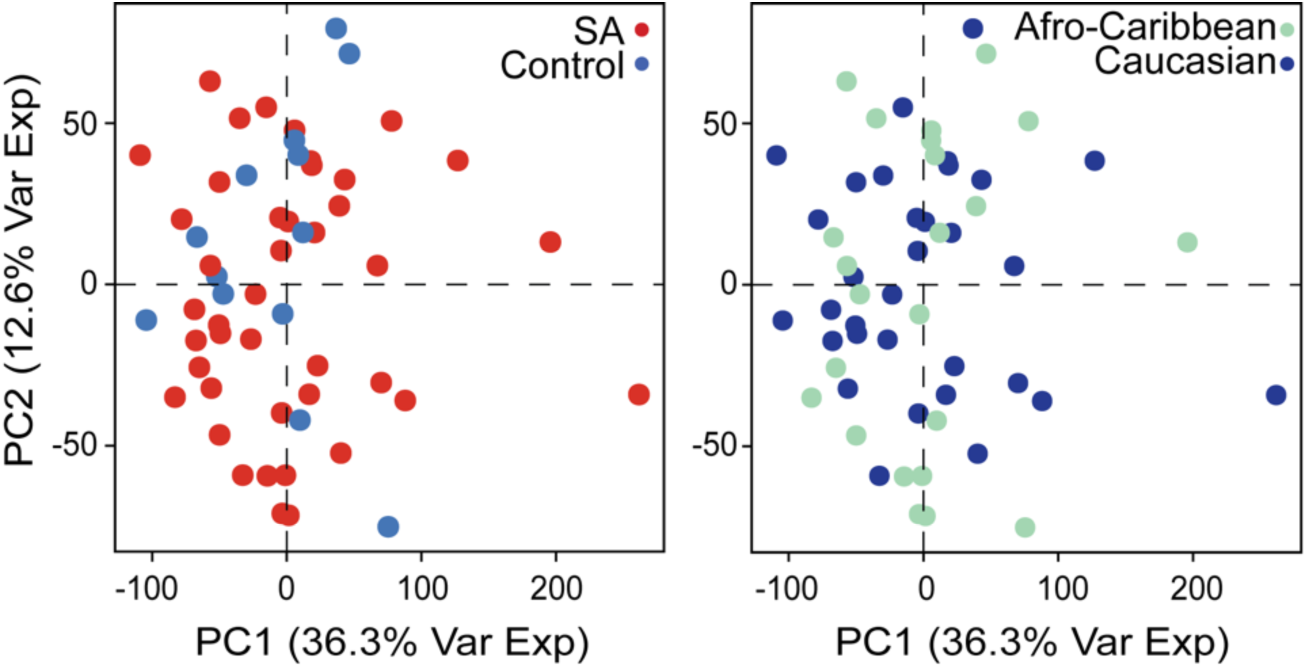
Principal component analysis of the merged Afro-Caribbean and Caucasian dataset. Human patients are labeled by their disease condition and their association with a demographic cohort (SA=sarcopenia).

**Supplementary Figure 3.**
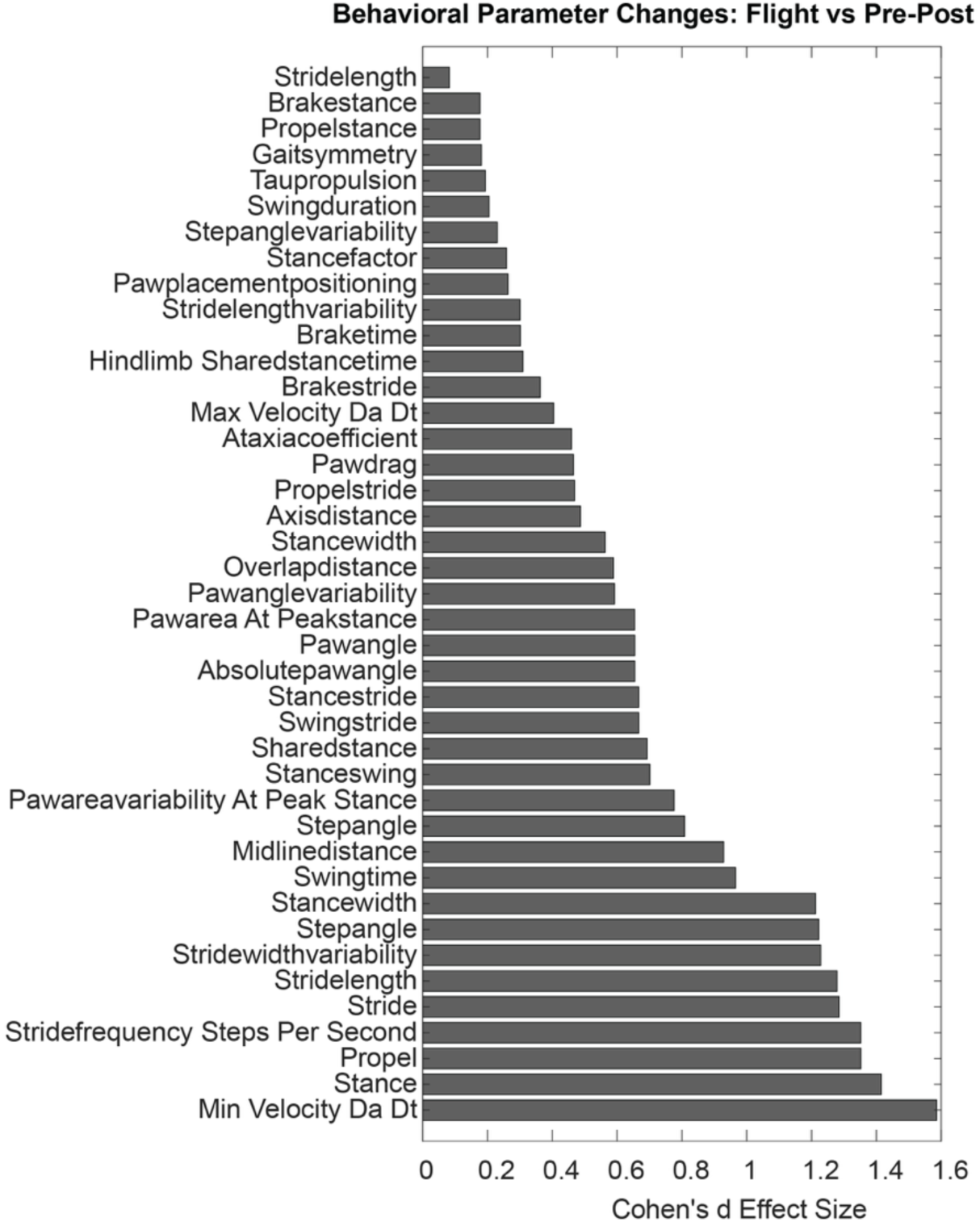
Effect size of behaviors pre- and post-flight. All gait parameter effect sizes were calculated between pre- and post-flight time points.

